# Genetic manipulation using hepatocyte-targeting adeno-associated viral vectors has minimal off-target effects

**DOI:** 10.1101/2021.02.19.431990

**Authors:** Christos Kiourtis, Ania Wilczynska, Colin Nixon, William Clark, Stephanie May, Thomas G Bird

## Abstract

Mice are a widely used pre-clinical model system in large part due to their potential for genetic manipulation. The ability to manipulate gene expression in specific cells under temporal control is a powerful experimental tool. The liver is central to metabolic homeostasis and a site of many diseases, making the targeting of hepatocytes attractive. Adeno-Associated Virus 8 (AAV8) vectors are valuable instruments for the manipulation of hepatocellular gene expression. However, their off-target effects in mice have not been thoroughly explored. Here, we sought to identify the short-term off-target effects of AAV8 administration in mice. To do this, we injected C57BL/6J Wild-Type mice with either recombinant AAV8 vectors expressing Cre recombinase or empty AAV8 vectors and characterised the changes in general health and in liver physiology, histology and transcriptomics compared to uninjected controls over 1 week. We observed an acute and transient reduction in homeostatic liver proliferation together with induction of the DNA damage marker γH2AX following AAV8 administration. The latter was enhanced upon Cre recombinase expression by the vector. Furthermore, we observed transcriptional changes in genes involved in circadian rhythm and response to infection. Notably, there were no additional transcriptomic changes upon expression of Cre recombinase by the AAV8 vector. Overall, there was no evidence of liver injury, dysfunction or leukocyte infiltration following AAV8 infection. These data support the use of AAV8-based Cre recombinase delivery as a specific tool for hepatocellular gene manipulation with minimal effects on murine physiology but highlight the off target effects of these systems.

**Summary statement:** This paper provides a comprehensive characterisation of the short-term effects of administration of Adeno-Associated Virus 8 on murine physiology, liver histology and liver transcriptome.

## Introduction

Animal models have improved our understanding and therapies for human disease. The mouse is a prototypical model organism that is widely used for a number of reasons, including its similarities with human physiology, breeding efficiency and ease of handling, cost efficiency and the range of available genetic models. Due to the latter particularly, mice have become the most widely used *in vivo* pre-clinical model system (Rosenthal and Brown, 2007). Manipulation of gene expression in this model organism has come a long way from whole body knock-out (KO) to the current point that we are able to introduce point mutations in a tissue specific manner through CRISPR-Cas9 genomic editing (Sauer and Henderson, 1988; Wilson, 1996; Lee, Yoon and Kim, 2020; Lundin *et al*.,2020). The Cre-Lox system, although less flexible compared to CRISPR, remains widely used for the manipulation of gene expression in mice and is an readily applicable means of genomic editing with high reproducibility.

Taking advantage of the Cre-Lox system, Adeno-Associated Viruses (AAVs) are an important vector system for gene expression manipulation and their use has risen dramatically in the last 20 years. AAVs, being replication deficient, are a relatively safe and efficient way to express the Cre recombinase, overexpress specific proteins or introduce shRNA into *in vivo* model systems. AAVs are small (20nm), single-stranded DNA viruses that belong to the family of Parvoviridae. They elicit a very mild immune response, especially the recombinant AAV vectors (rAAVs) which have undergone modifications to partly evade the immune system (Rogers *et al*.,2011; Rabinowitz, Chan and Samulski, 2019). There are different serotypes of AAV (AAV1, 2, 4, 5, 6, 7, 8, and 9), each of which exhibits a different transduction efficiency in the different target tissues (Zincarelli *et al*.,2008). In mice, after infecting their target cells, AAVs enter the cell nucleus where they persist in an episomal form and only rarely integrate into the host genome (Duan *et al*.,1999; Miller, Petek and Russell, 2004).

The liver is the largest solid organ in the body and is a frequent site of organ-specific and systemic diseases and a frequent site of tumour metastasis. In liver biology, studying hepatocytes is particularly important as they constitute the majority of liver cells, comprising around 60% of total liver mass. Hepatocytes perform most of the synthetic and detoxification functions of the liver and are responsible for liver regeneration as well as being the cell of origin of the majority of primary liver cancers (Müller, Bird and Nault, 2020). As a result, genetic manipulation of hepatocytes is a powerful tool in the study of liver disease.

There are a number of ways to manipulate hepatocellular gene expression (Kellendonk *et al*.,2000). Currently, a widely used approach is to target hepatocytes with an AAV-based vector. rAAV8 is a commonly used AAV serotype due to its strong propensity to transduce hepatocytes (Nakai *et al*.,2005). rAAV8-mediated hepatocellular gene editing has multiple applications including gene therapy (Smith *et al*.,2011), lineage tracing experiments, gene deletion or gene overexpression in all or specific populations of the hepatocytes. Through the insertion of tissue specific promoters vector tropism for a specific tissue or cell type can be enhanced. In particular, the Cre recombinase together with a hepatocyte-specific promoter like the Thyroxin Binding Globulin (*TBG*) promoter can be incorporated into the AAV8 genome and this is reported to be a specific means of Cre recombinase expression in hepatocytes, while avoiding undesired transduction of extrahepatic cells (Nakai *et al*.,2005; Malato *et al*.,2011; Lee *et al*.,2020). The number of transduced hepatocytes is proportional to the dose (i.e. genetic copies) of AAV8-*TBG* vector that are administered; the higher the dose of the vector, the more hepatocytes will be transduced. This allows the study of deleting/overexpressing a gene in the whole liver parenchyma (Bird *et al*.,2018) or in a small number of hepatocytes using comparatively fewer genetic copies of vector. Alternatively, instead of the Cre recombinase, it is possible to deliver other constructs as “cargo” (e.g. expression of shRNAs or ectopic proteins) to hepatocytes using this approach; for example, administration of the AAV8-*TBG-*P21 vector results in P21 upregulation in hepatocytes, inhibiting their ability to proliferate (Raven *et al*.,2017). Expression of ectopic proteins with AAV vectors has been reported to last for several months, at least in post-mitotic cells (Duan *et al*.,1999).

The AAV8 system theoretically allows for manipulation of gene expression at a desired time point and without inducing toxicity or the risk of genetic ‘leakiness’ through an endogenous Cre allele. This is in comparison to other models like the Albumin-Cre mice, where the Cre recombinase is constitutively expressed from embryonic life and is therefore not temporally controlled, or tamoxifen-mediated manipulation of gene expression, where tamoxifen has been reported to induce toxicity (Gao *et al*.,2016; Keeley, Horita and Samuelson, 2019). As such, AAV8-TBG is widely used in order to recombine the majority of the hepatocytes (90-95%) and study the effects of gene expression changes in the whole liver serving as a single hit, hepatocyte-specific gene knock-out/overexpression.

With the report that AAVs may have long lasting effects upon the liver epithelium, including rare cancers, it is clear than transduction with AAV is not entirely benign (Nault *et al*.,2015). Even though in humans evidence suggests that the immune system might compromise AAV8 efficiency (partly due to cross-immunity with Adenoviruses) there haven’t been detailed studies on the murine immune response against AAV8 (Boutin *et al*.,2010; Mendell *et al*.,2010; Calcedo *et al*.,2011). Furthermore, as rAAV8 rarely integrates into the murine host genome, it seems unlikely that it would cause significant genotoxicity. In one study investigating the long term effects of AAV2-hFIX16 (which results in liver-specific expression of clotting factor IX) in liver tumourigenesis in mice, it was found that there was no association between tissue from hepatocellular carcinomas (HCCs) and AAV copy numbers (Li, Malani and Hamilton, 2011).

Transcriptome-wide studies are commonly performed on whole liver lysates or isolated liver cell fractions of mice treated with AAV8-*TBG*-Cre. These transcriptomics analyses can give valuable information on the effects following manipulation of hepatocellular gene expression via AAV8-*TBG*-Cre. However, a potential effect on the transcriptome by the AAV8 vector or by its “cargo” (i.e. the Cre recombinase or other protein expressed by the vector) should be taken into consideration when performing and interpreting such studies. To our knowledge there are currently no studies addressing whether AAV vectors (and in particular AAV8-*TBG*) alone have an effect on the liver transcriptome.

Overall, there is a lack of descriptive studies on the effects of systemic AAV8 administration in mice. Therefore, to address this shortfall we investigated the short-term off-target effects of systemic AAV8-TBG administration in Wild-Type (WT) mice. After intravenous (i.v.) injection of AAV8-*TBG*-Cre (expressing Cre recombinase) or AAV8-*TBG*-Null (expressing a scrambled sequence) at dosing resulting in transduction across the entire hepatocyte compartment we examined both liver specific and systemic alterations in WT mice. Using blood analysis combined with immunohistochemistry and transcriptomics analysis we describe the effects occurring over a week post transduction. These data confirm minor off target effects following transduction using this experimental strategy and serve as a reference tool for the research community.

## Results

### AAV8-*TBG* is hepatocyte-specific

We first examined the tissue and cell specificity of AAV8-TBG using mice homozygote for the R26-LSL-tdTomato allele on a C57BL/6 background by simultaneous injection with AAV8-*TBG*-Cre and AAV8-*TBG*-GFP (herein referred to as AAV-Cre and AAV-GFP respectively) (Fig. 1A). The cells expressing the GFP and RFP reporters 7 days after AAV8 injection were assessed histologically first in the liver, demonstrating that almost all hepatocytes expressed the reporters (Fig. 1B), consistent with previous reports using this (Bird *et al*.,2018; Gay *et al*.,2019) and other AAV8-Cre constructs (Malato *et al*.,2011). There was no evidence of recombination of biliary epithelium or other non-parenchymal populations in the liver. Interestingly, while RFP staining was distributed evenly across the hepatocytes, the GFP distribution was more irregular and its intensity varied among hepatocytes, with a tendency for more intense staining in the hepatocytes surrounding the central vein (pericentral hepatocytes of Zone 3) (Fig. 1B). Notably, when we checked for reporter expression in other organs, we observed labelling of very few cells in the duodenum, kidney, pancreas, lung and the spleen (Fig. 1C, 1D). The apparent GFP positivity observed in the duodenum and the spleen of uninjected mice (Fig. 1C, inset images) appears as non-specific background staining. These data show, in agreement with other studies (Wang *et al*.,2010; Bell *et al*.,2011), that AAV8-*TBG*-mediated gene targeting is highly specific for hepatocytes with negligible targeting of extra-hepatic tissues.

**Figure 1:**
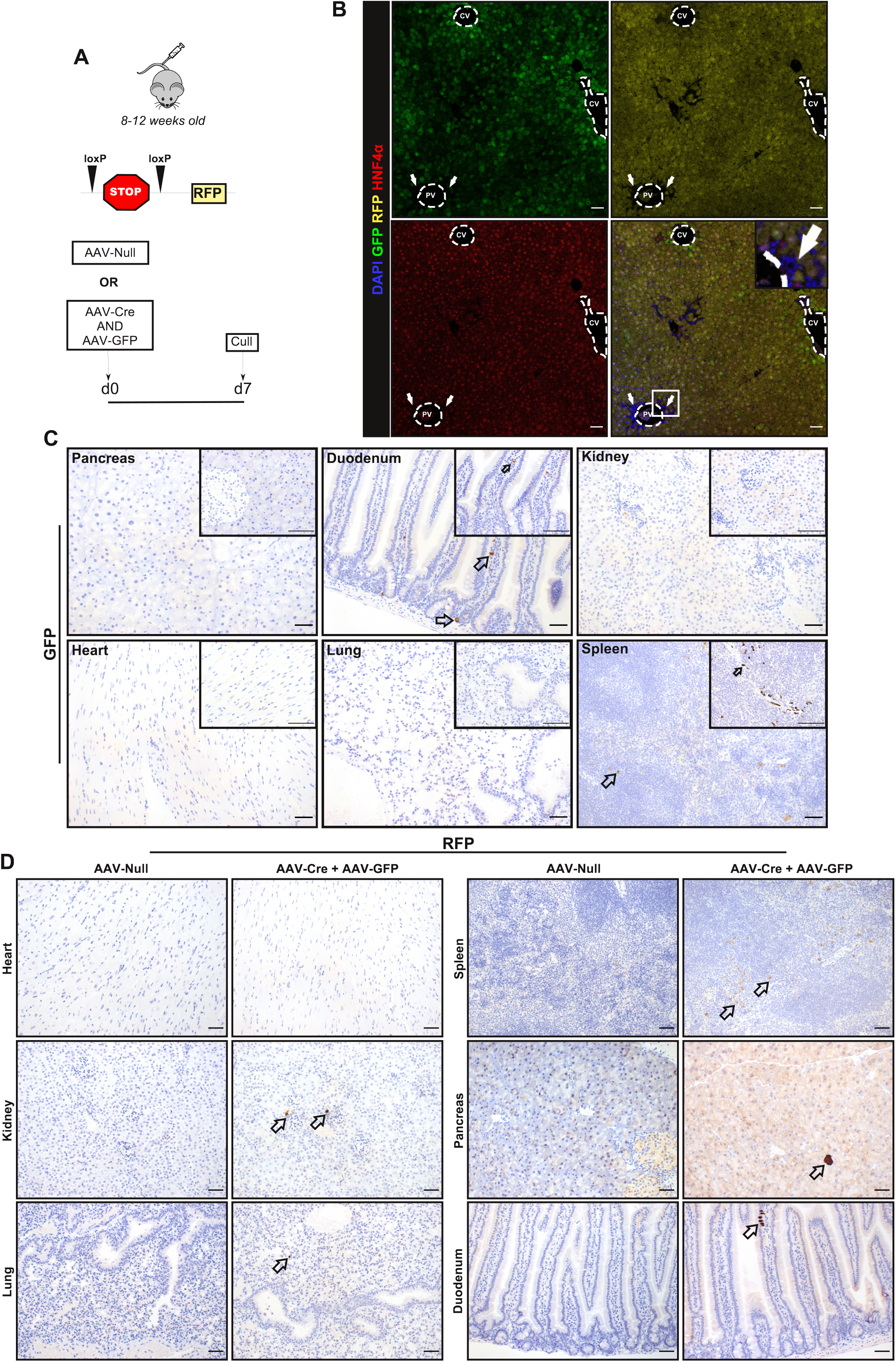
AAV8-*TBG* vectors specifically target the hepatocytes. **(A**) Schematic of the experimental design; 8-12 week old male LSL-RFP mice on a C57BL/6 background (n=6) were i.v. injected with AAV-Cre and AAV-GFP at the same dose (2×10^11^ GC/mouse). LSL-RFP mice (n=4) injected with AAV-Null served as controls. 7 days later their livers were harvested for analysis. **(B)** Representative images from liver sections stained for DAPI (blue), GFP (green), RFP (yellow) and the hepatocyte-specific marker HNF4α (magenta), showing the hepatocellular specificity of the AAV8-*TBG* vectors. Arrows highlight the unlabelled bile ducts. CV = Central Vein; PV = Portal Vein. **(C)** Representative images of GFP immunohistochemistry in the pancreas, duodenum, kidney, heart, lung and spleen of mice injected with AAV-Cre and AAV-GFP. The inset images are from GFP-stained liver sections from uninjected WT mice (i.e. mice not injected with either AAV-Cre or AAV-GFP, representative images from n=3 mice). Arrows highlight GFP-positive cells **(D)** Immunohistochemistry for RFP in the kidney, pancreas, spleen, heart, lung and duodenum of the mice described in 1A. Arrows highlight RFP-positive cells. All scale bars are 50μm.

### Systemic administration of AAV8-*TBG* does not affect the general health of mice

To investigate the off-target effects of systemic AAV8-*TBG* administration, WT mice were i.v. injected with AAV8-*TBG*-Null (herein referred to as AAV-Null) or AAV-Cre. Mice were then culled 2, 4 or 7 days post AAV8-*TBG* injection and compared to uninjected controls using a number of clinical parameters (Fig. 2A). Starting at a similar body weight at day 0 (Fig. S1A), the mice showed no significant changes in body weight and gradually gained weight at a normal rate for their age during the week after AAV-Null or AAV-Cre, regardless of the group (Fig. 2B). Haematology analysis showed no changes in haematocrit, red blood cells and platelets (Fig. 2C). Reflecting the reported mild inflammatory response elicited by AAVs, we did not observe significant changes in circulating total white blood cells, monocytes and neutrophils (Fig. 2D, 2E, Fig. S1B). We observed a significant difference in circulating lymphocytes between AAV-Null day 4 and AAV-Cre day 7 groups (Fig. 2E). This did not translate to a significant change in the relative numbers of lymphocytes in these groups (Fig. S1B). Overall, we did not observe any impact on general health of mice a week after AAV-Null or AAV-Cre administration.

**Figure 2:**
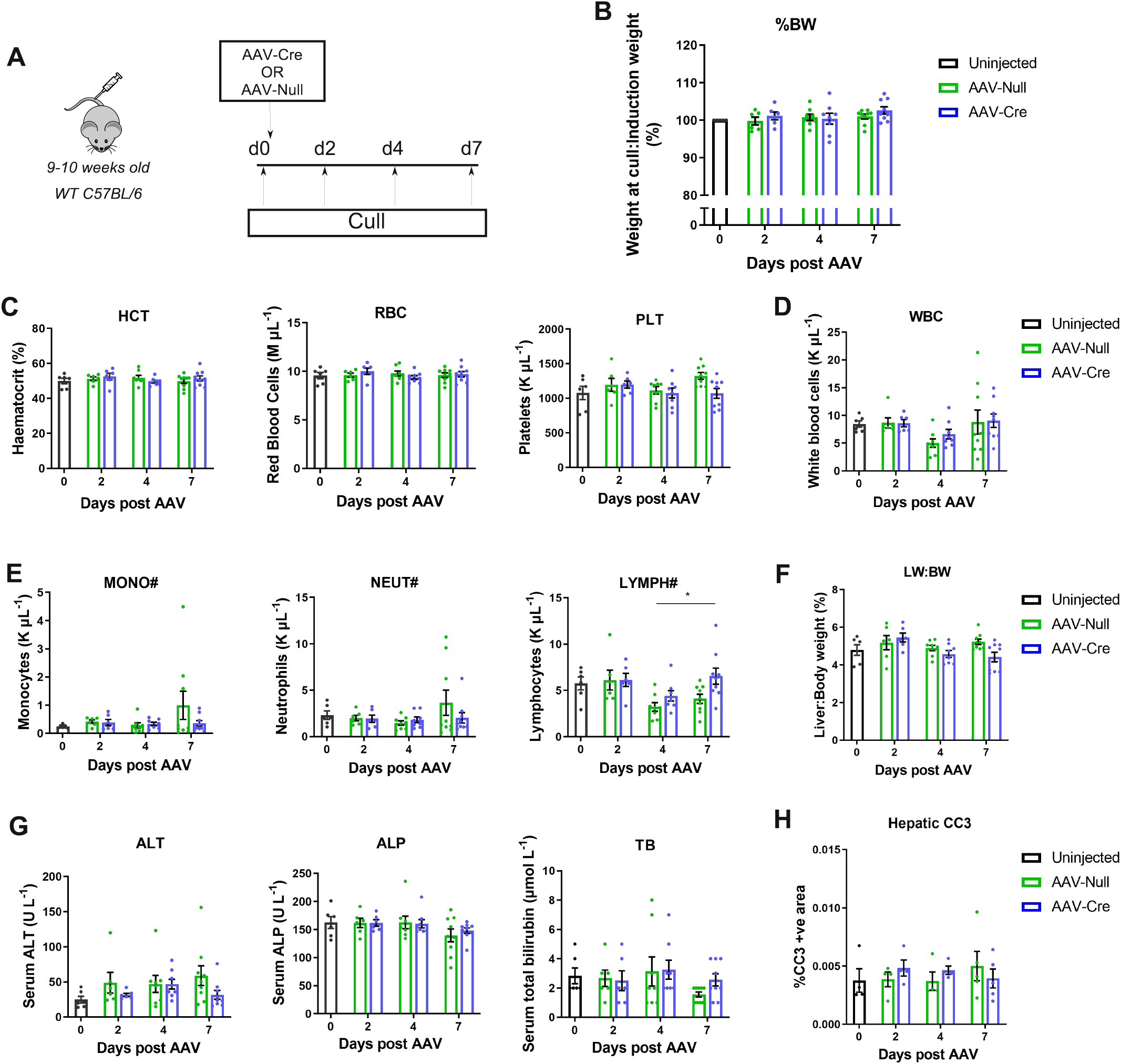
Systemic administration of AAV8-*TBG* has minimal effects on general health causing neither liver injury nor impaired liver function. **(A)** Schematic of experimental outline. Male C57BL/6J WT mice (n=46) were injected i.v. with either AAV-Null or AAV-Cre. Uninjected control mice (n=6) from the same stock were culled on the day that the rest of the mice were injected with AAV8-*TBG* (day 0). The injected mice were culled 2 (n=12; 6 AAV-Null and 6 AAV-Cre), 4 (n=16; 8 AAV-Null and 8 AAV-Cre) or 7 (n=18; 9 AAV-Null and 9 AAV-Cre) days after injection. **(B)** Body weight (BW) at cull in relation to body weight at day 0 for the mice described in 2A (each data point represents 1 mouse). Kruskal-Wallis test showed no statistically significant differences. **(C)** Haematocrit (HCT), Red Blood Cell (RBC) and Platelet (PLT) counts for uninjected, AAV-Null and AAV-Cre mice. One-way ANOVA showed no statistically significant differences. **(D)** Circulating White Blood Cell (WBC) counts for uninjected, AAV-Null and AAV-Cre mice. Kruskal-Wallis test. **(E)** Absolute blood counts of circulating Neutrophils (NEUT#), Monocytes (MONO#) and Lymphocytes (LYMPH#) for uninjected, AAV-Null and AAV-Cre mice. P= *<0.05, one-way ANOVA. **(F)** Liver weight to body weight ratio (LW:BW) of uninjected, AAV-Null and AAV-Cre mice. One-way ANOVA showed no statistically significant differences. **(G)** Alanine aminotransferase (ALT), Alkaline phosphatase (ALP) and Total Bilirubin (TB) in the plasma of uninjected, AAV-Null and AAV-Cre mice. One-way ANOVA (for ALT) or Kruskal-Wallis test (for ALP and Bilirubin) showed no statistically significant differences. **(H)** Area quantification for Cleaved Caspase 3 (CC3) and P21 (representative images in Fig. S2). Kruskal-Wallis test showed no statistically significant differences. The bars on all graphs are mean ± S.E.M.

### AAV8-*TBG* vectors do not cause liver damage and do not affect liver function

Next, having demonstrated hepatocyte-specific targeting, we proceeded to assess the effects of AAV8-*TBG* on the liver specifically. Livers were normal macroscopically and we did not observe any changes in liver size or liver histology microscopiocally (as assessed by H&E staining) in response to AAV8 (Fig. 2F, S1C, S2). Similarly, serum levels of Alanine aminotransferase (ALT) and Alkaline phosphatase (ALP) (markers of liver necrosis and bile duct damage respectively) remained at baseline levels at every time point (Fig. 2G). Assessing liver function, serum bilirubin levels also remained unaffected as did serum levels of Total protein, Albumin, Globulin and Albumin:Globulin ratio (Fig 2G, S1D). Examining hepatic cell death in more detail, we performed immunohistochemistry for the apoptosis-specific marker Cleaved Caspase 3 (CC3). No changes in apoptotic cell death were observed at any time point (Fig. 2H). There was no change in serum urea levels, however creatinine was significantly increased in AAV-Null day 4 mice, returning to baseline at day 7 (Fig. S1E). Therefore, we found no evidence of liver damage or dysfunction after AAV8-*TBG* administration during the times when transduction and generic recombination occur.

We next examined intrahepatic leukocyte populations to test whether a demonstrable local immune response occurred in the liver. Using the pan-leukocyte marker CD45, we didn’t observe any change in overall leukocyte numbers or distribution (Fig. 3A, S2). The use of more specific leukocyte markers for neutrophils (Ly6G), macrophages (F4/80), T-cells (CD3) and B-cells (B220) also demonstrated no significant differences in these populations either in number or distribution at any timepoint (Fig. 3A, S2, S3). Therefore we find no evidence of histological inflammation or inflammatory response to biologically relevant AAV8 dosing.

**Figure 3:**
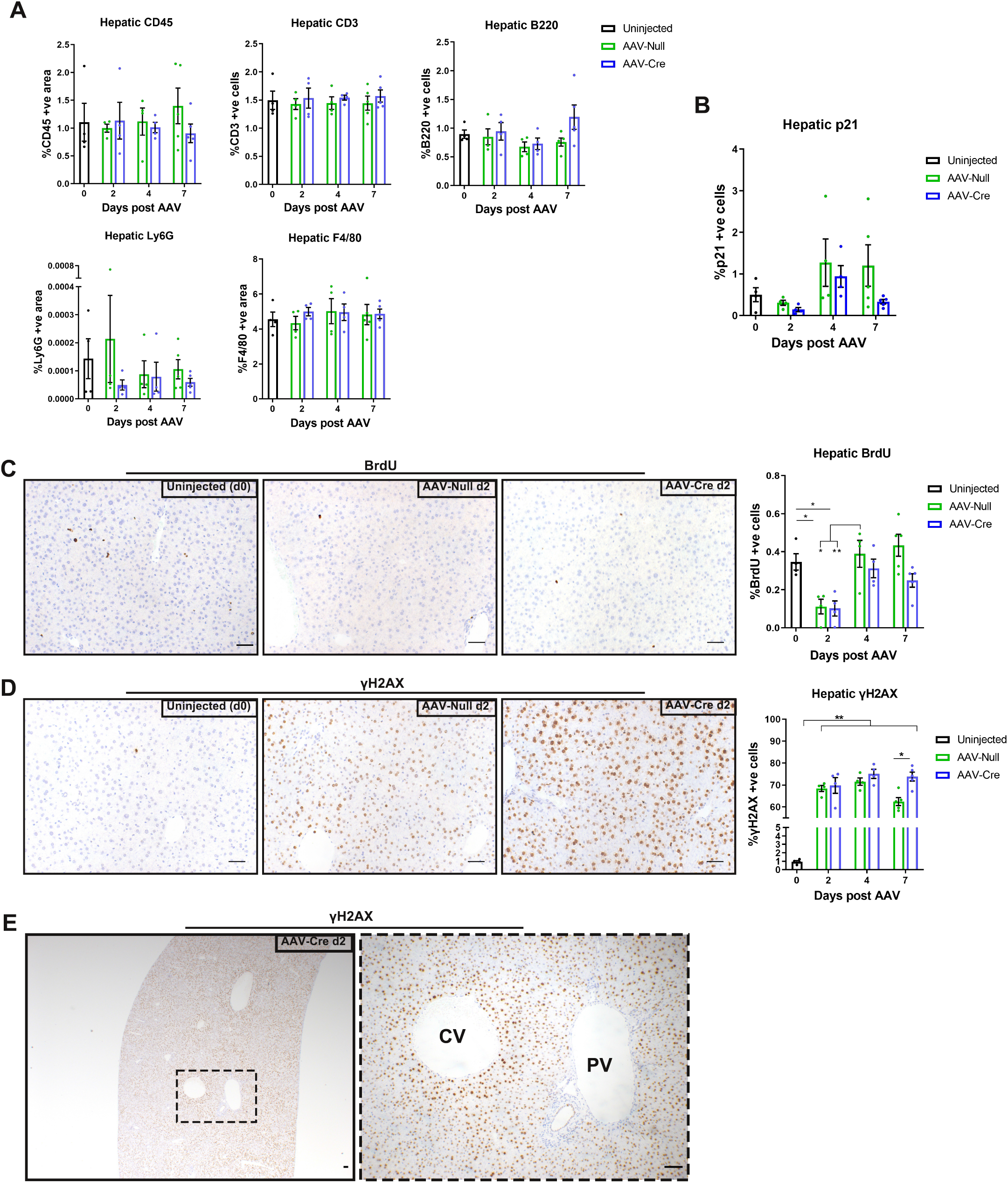
AAV8-*TBG* vectors affect the hepatocellular cell cycle and result in a DNA damage response. **(A)** Quantification of hepatic CD45, CD3, B220, F4/80 and Ly6G based on positive area/total liver area (CD45, F4/80, Ly6G) or positive cells as a percentage of total cells (CD3, B220) after immunohistochemical detection (representative images for each time point in Fig. S2, S3). One-way ANOVA (CD3), Kruskal-Wallis test (CD45 and Ly6G) or Brown-Forsythe and Welch ANOVA (B220 and F4/80) showed no statistically significant differences. **(B)** Quantification of hepatic P21 positive cells presented after immunohistochemical detection (representative images for each time point in Fig. S4). Brown-Forsythe and Welch ANOVA showed no statistically significant differences. **(C)** Quantification of liver cells positive for the S-phase marker BrdU and representative immunohistochemistry images (additional images for each time point are shown in Fig. S4); one-way ANOVA; P= *<0.05, **<0.01. **(D)** Quantification of γH2AX positive liver cells and representative immunohistochemistry images (additional images for each time point are shown in Fig. S4); Brown-Forsythe and Welch ANOVA; P= *<0.05, **<0.01. **(E)** Representative liver section stained for γH2AX showing zonal staining particularly in the pericentral area (Zone 3). CV = Central vein, PV = Portal vein. For all graphs n=4 in all groups apart from AAV-Null d7 and AAV-Cre d7 where n=5. The bars on all graphs are mean ± S.E.M and all scale bars are 50μm.

### AAV8-*TBG* vectors affect the cell cycle of liver cells and induce expression of the DNA damage marker γH2AX in the liver

Viral infection of mammalian cells is, through a variety of well characterised mechanisms, known to affect several cellular processes including cell cycle, DNA damage response and the release of Damage-Associated Molecular Patterns (DAMPs) (Loo and Gale, 2011; Dou *et al*.,2017; Motwani, Pesiridis and Fitzgerald, 2019). To address whether AAV8-*TBG* vectors can induce such changes, we first stained liver sections for the cell cycle inhibitor *Cdkn1a* (P21) or for BrdU to determine changes in the cell cycle status of liver cells. Whilst there was no significant change in hepatic P21 at any timepoint in either group, there was a significant transient reduction of BrdU positive cells at day 2 post AAV8-*TBG* administration (Fig. 3B, 3C, S4). Next, we assessed the presence and extent of hepatic DNA damage by staining liver sections for the DNA damage marker γH2AX. We observed a marked increase in γH2AX at day 2, persisting until day 7, both in the AAV-Null and in the AAV-Cre groups (Fig. 3D, S4). Moreover, treatment with AAV-Cre resulted in a stronger γH2AX response (Fig. S1F, S4). Notably, γH2AX staining was stronger in the pericentral hepatocytes (Fig. 3E). Our data reveal an acute but transient reduction in hepatic proliferation as well as an increase in hepatic γH2AX following systemic AAV8 administration.

### AAV8-*TBG* vectors induce circadian rhythm-and infection-related transcriptional changes

As a broader and unbiased assessment of AAV8-*TBG* vectors effects we next explored their effect on the liver transcriptome by performing RNA-seq on whole liver lysates from our AAV8-*TBG*-treated and uninjected control mice (Fig. 4A). In general, there was a strong degree of similarity among all samples by Principal Component Analysis (PCA) (Fig. 4B). We interrogated this transcriptomics data in more detail, starting with the AAV8-*TBG* cargo in each group. Here we observed that there was a gradual increase in the number of the respective AAV8-*TBG* transcripts detected from day 2 to day 7 (Fig. 4C). Transcript number was also influenced by the specific cargo; expression of Cre transcript was lower than that of the transcript expressed by AAV-Null. Next, we performed pathway analysis in order to identify global transcriptional changes. This revealed two broad transcriptional programmes that were altered among the different timepoints; immune response-related changes and circadian rhythm changes (Fig. 4D). Notably, using this unbiased approach we did not observe any transcriptional changes associated with DNA damage response.

**Figure 4:**
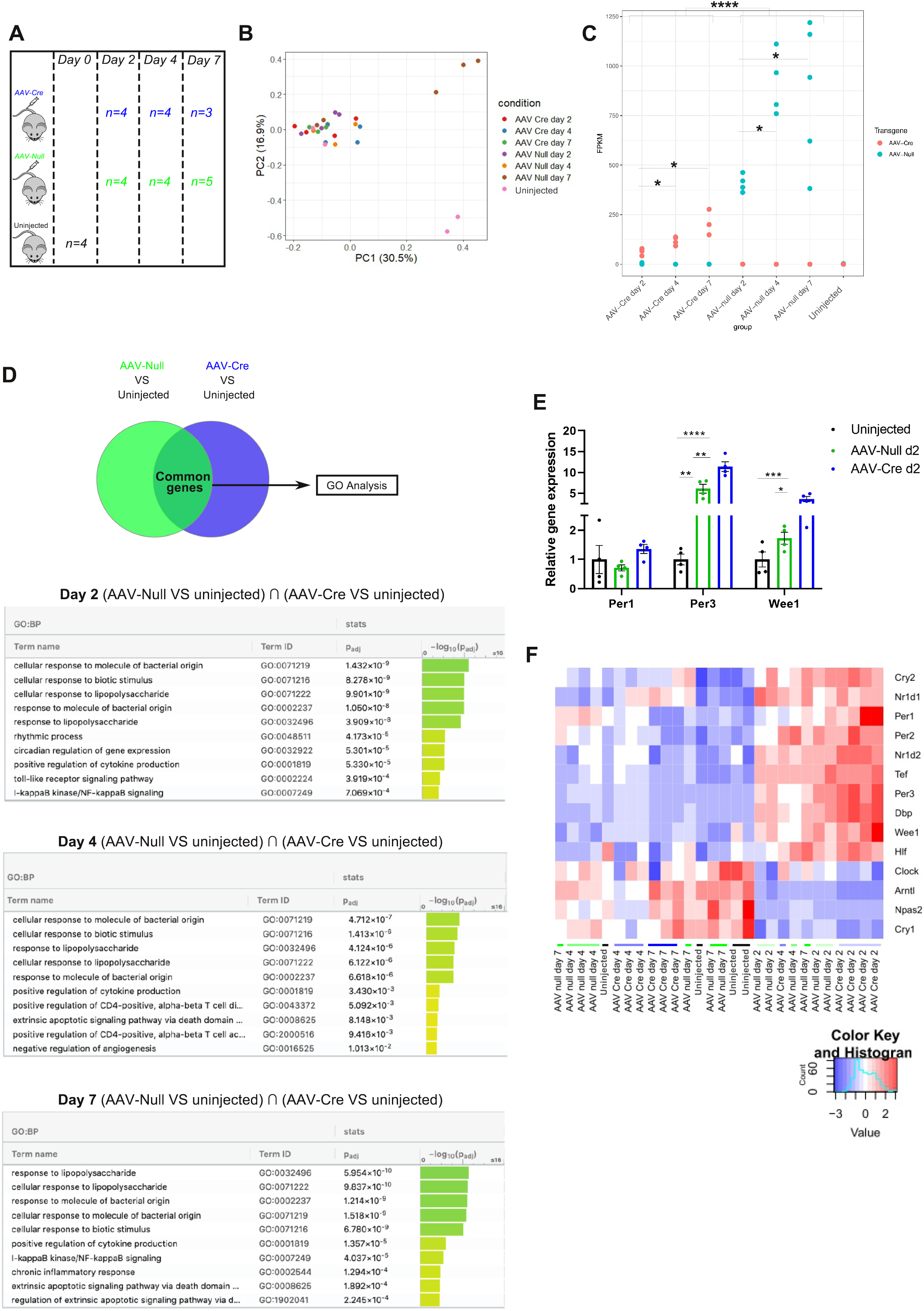
Short-term temporal effects of AAV8-TBG upon the liver transcriptome. **(A)** Schematic of the samples used for RNA-seq. Whole liver lysates from 4 uninjected, 13 AAV-Null (n=4 at day 2, n=4 at day 4 and n=5 at day 7 post injection) and 11 AAV-Cre (n=4 at day 2, n=4 at day 4 and n=3 at day 7 post injection) mice were used. **(B)** Principal Component Analysis (PCA) plot of the samples used for RNA-seq. **(C)** Quantity of the transcripts encoded by AAV-Cre (sequence of the Cre recombinase) or AAV-Null (scrambled sequence) in the different conditions represented as Fragments Per Kilobase of transcript per Million mapped reads (FPKMs). 2-way ANOVA. *P < 0.05; ****P < 0.0001. **(D)** Gene Ontology (GO) analysis comparing the differentially expressed genes shared between AAV-Null and AAV-Cre mice after each group is compared to uninjected mice (AAV-Null VS uninjected ∩ AAV-Cre VS uninjected) mice at day 2, 4 and 7. **(E)** RT-qPCR for Per1, Per3 and Wee1. Fold change expression was calculated by normalizing to the uninjected mice for each gene. n=4 for each group. Kruskal-Wallis test (Per1) or one-way ANOVA (Per3, Wee1). *P < 0.05; **P < 0.01; ***P < 0.001 and ****P < 0.0001. The bars are mean ± S.E.M. **(F)** Unsupervised heatmap showing the differential expression of major genes involved in circadian rhythm regulation.

Having observed prominent effects on cellular proliferation at day 2, we focused on the circadian rhythm process that was specific for this timepoint. First, we validated the expression of specific genes involved in circadian rhythm (Takahashi, 2017) observing similar trends of expression to those of the RNA-seq (Fig. 4E, F). Similarly to the reduced proliferation at day 2, the changes in circadian rhythm were viral-specific rather than cargo-specific; the change was observed at a specific time point regardless of the cargo (Fig. 4F). Furthermore, some of the genes involved in these networks (Wee1, Tef) have been described to regulate cell cycle (Russell and Nurse, 1987; Rowley, Hudson and Young, 1992; Yang *et al*.,2019). Overall, our transcriptomic data reveals changes in genes involved in the circadian rhythm as well as in inflammation and immunity.

## Discussion

AAV8-*TBG* vectors are an established means for hepatocyte-specific manipulation of gene expression *in vivo*. In this study we show that AAV8-*TBG* vectors have both a high degree of specificity and minimal off-target effects. Therefore, they serve as a reliable and efficient experimental tool. To our knowledge, our study is the first one to systematically examine these effects in the liver of WT mice. We demonstrate that mouse health is generally unaffected by AAV8-*TBG* vectors as the body and liver weights exhibited the expected growth. No inflammatory response, either systemic or intrahepatic, was observed and liver histology and function remained normal. However, we have identified some subtle phenotypes that are induced by AAV8-*TBG* vectors, which should be taken into account when using this system for *in vivo* experiments in mice. These observations highlight that AAV8-*TBG* vectors are not entirely benign.

The specific targeting of hepatocytes was demonstrated by 2 reporters, RFP and GFP. Importantly, even though there were a few labelled cells in extra-hepatic tissues in our study, AAV8-*TBG* vectors showed highly specific tropism for hepatocytes as previously reported (Wang *et al*.,2010; Bell *et al*.,2011). When considering phenotypic modification of cells, a low level of off-target recombination is unlikely to significantly affect short term studies, however it should be considered particularly when performing longer term experiments where modified cells may expand clonally.

We note differences in the labelling pattern between the 2 reporters; RFP evenly labelled almost all of the hepatocytes, while fluorescent intensity of GFP was more heterogeneous across zones, showing preference for the pericentral hepatocytes (Zone 3), but also among cells within the same zone. We suggest that this is explained by the different mechanisms of labelling. Expression of the tdTomato gene is endogenously regulated and protein expression depends on recombination following Cre expression by the AAV8-*TBG* vector; once Cre is expressed and the LSL cassette excised, there is continuous expression of RFP protein by the host genome. On the other hand, GFP is expressed directly from the AAV8-*TBG* vector. Therefore, its expression is predicted to vary from cell to cell depending on the quantity of viral copies delivered to each cell. The preferential labelling of pericentral hepatocytes by AAV8-*TBG*-GFP in mice has been demonstrated by others (Wang *et al*.,2010; Bell *et al*.,2011) but the exact mechanism remains unclear. It has been reported that a stronger “pericentral tropism” of AAV8 may underlie this (Bell *et al*.,2011), rather than differential expression of *TBG* across the liver zones. This effect was also apparent by the zonal distribution of γH2AX positivity. Here we also observed zonal differences which are further exacerbated by the expression of Cre recombinase, further supporting a zonal preponderance for higher tropism/expression of cargo in pericentral hepatocytes.

One of the key findings of this study is the widespread DNA damage response observed in the liver, as manifested by the increase in γH2AX. It has been previously shown that AAVs can, upon infection, induce DNA damage and mobilize the DNA repair machinery of the host cell in order to achieve the circular episomal form in which AAVs persist in the host cell (Schwartz *et al*.,2009; Cataldi and McCarty, 2013). These studies, mostly performed *in vitro*, identify DNA-PKcs as a key mediator of this process, with γH2AX being one of the DNA damage response components involved. Our study confirms the increase of hepatocellular γH2AX in mice *in vivo* in response to AAV-Null infection. The enhanced DNA damage response observed in the mice injected with AAV-Cre could be explained by additional, non-specific DNA damage induced by the Cre recombinase. This enzyme can unselectively cut DNA at non-Lox sites (Loonstra *et al*.,2001; Janbandhu, Moik and Fässler, 2014; Pépin *et al*.,2016; Lam *et al*.,2019). Lastly, it is important to highlight that, in our study, in spite of the increase in hepatic γH2AX, there were no apparent changes in histology or gene expression related to DNA damage.

The observed decrease of proliferation on day 2 in both AAV-Null and AAV-Cre indicates that this is an AAV8-*TBG* mediated effect rather than solely one mediated by the Cre recombinase as has been described by others (Loonstra *et al*.,2001). This reduction of proliferation is unlikely to be biologically significant in the longer term as it affects a small proportion of liver cells (a drop of 0.2 percentile units compared to uninjected controls). Nonetheless, it is possible that the affected liver cells are important for specific functions, so further characterisation of this phenotype should be considered depending on the experimental question being tested. One transcriptional process that was altered in AAV8-*TBG*-treated mice was the circadian rhythm, with the changes taking place on day 2. Circadian rhythm is classically viewed as an internal biological clock manifested by oscillations in gene expression and which is mainly affected by photoperiodism. The liver however has an additional autonomous internal clock and thus it is not entirely dependent on photoperiodism (Koronowski *et al*.,2019; Li *et al*.,2020). Our transcriptomics analysis identified several genes involved in circadian rhythm that are differentially expressed at day 2. As some of these genes have been implicated in the control of cell cycle (Matsuo *et al*.,2003; Zhou *et al*.,2018), It is possible that these transcriptional changes are related to the mild decrease in hepatic proliferation we observed at day 2.

Our transcriptomics analysis of whole liver lysates revealed that AAV8-*TBG* vectors can induce transcriptional changes in the liver. The most prominent transcriptional changes identified in GO analysis are related to infection and inflammation processes and were observed in all the time points of the study. Given the viral nature of AAV8-*TBG* vectors, it is perhaps unsurprising to observe these transcriptional responses in the infected cells. However, in our hands, this transcriptional response to infection did not result in a demonstrable modification of the numbers of immune cells as a marker of inflammatory response. Nevertheless, these transcriptional changes should be considered in experiments with AAV8-*TBG*, especially when the focus of the study is related to the immune system and/or inflammation.

One limitation of our work is that we have not explored the longer term consequences of AAV8 use in WT animals. We have observed long term hepatic expression of GFP following AAV8-*TBG*-GFP administration to animals for over 200 days (Valentin Barthet, personal communication). Persistent expression of AAV8-*TBG*-driven GFP in the liver suggests persistence of AAV8-*TBG* vectors in the hepatocytes. Therefore, it would be interesting to characterise the long term effects of AAV8-*TBG* vectors in mice.

In this study we describe the short term off-target effects of systemic administration of AAV8-*TBG* vectors in mice at a dose relevant for target delivery across the entire hepatocyte population. Although other studies have reported the some aspects of off-target effects of AAVs, these have mostly been performed *in vitro* and only explored specific hypothesis driven effects. In our study, the use of WT C57BL/6J mice to map the AAV8-*TBG* off-target effects, both systemic and liver-specific, makes our data relevant to that of other researchers. Additionally, the unbiased transcriptomics analysis serves to generally reassure about a lack of major off-target effects within hepatocytes when using this vector system, whilst acting as a useful tool for other researchers. In conclusion, our data show that AAV8-*TBG* vectors are a reliable and efficient tool for hepatocyte-specific genetic manipulation with minimal off-target effects.

## Materials and Methods

### Animal experiments

9-10 weeks old male C57BL/6J WT mice (*Mus musculus*) were purchased from Charles River UK. To minimise biological variability we obtained mice from as few litters as possible. The mice were housed in cages of 4-5 mice/cage in a licensed, specific pathogen-free environment facility under standard conditions with a 12 hr day/night cycle and ad libitum access to food and water. All experiments were carried out with ethical permission from the Animal Welfare and Ethical Review Body (AWERB) and in accordance with the ARRIVE guidelines (du Sert *et al*.,2020) and the Home Office guidelines (UK licence 70/8891; protocol 2).

AAV8 experimentation was performed as previously described (Bird *et al*.,2018). Briefly, stock AAV8.TBG.PI.Cre.rBG (AAV8-*TBG*-Cre) (Addgene, 107787-AAV8) or AAV8.TBG.PI.Null.bGH (AAV8-*TBG*-Null) (Addgene, 105536-AAV8) (stored at −80 °C) was thawed on ice, diluted in sterile PBS to achieve a working titre of 2×10^12^ genetic copies (GC)/ml and was subsequently stored at −20 °C until usage. On the day of the injection the diluted AAV was thawed and each mouse was injected via the tail vein with 100μl (2×10^11^ GC/mouse; mice in this study weighed from 22.4 – 29.4g at the time of injection). This dose has been previously shown to result in genetic recombination of nearly the total hepatocyte population (Bird *et al*.,2018). All mice were weighed on injection day (day 0) and on their respective cull day. Changes in body weight were compared to published data for this mouse strain (The Jackson Laboratory, Body Weight Chart #000664, URL (accessed on 26/11/2020): https://www.jax.org/jax-mice-and-services/strain-data-sheet-pages/body-weight-chart-000664#). The mice were sacrificed 2, 4 or 7 days post AAV8 administration. Male C57BL/6J mice from the same batch and of the same age without AAV8 administration (uninjected controls) served as baseline controls. All mice were culled between the hours of 11:00 and 15:00 on the day of harvest. All mice were injected with BrdU (Amersham, RPN201, 250μl per mouse) intraperitoneally 2 hrs before culling.

For the confirmation of tissue specificity of AAV8 we used 8-12 weeks old male mice on a C57BL/6 background that were homogygotes for the R26RLSL-tdTomato allele (LSL-RFP) (Madisen *et al*.,2010). These mice were injected on the same day with both AAV8-TBG-Cre and AAV8.TBG.PI.eGFP.WPRE.bGH (AAV8-*TBG*-GFP) (Addgene, 105535-AAV8), both at a dose of 2 x 10^11^ GC/mouse as described above. These mice were culled 7 days post AAV administration. LSL-RFP mice that were injected with 2 x 10^11^ GC of AAV8-TBG-Null and culled 7 days later served as controls for RFP expression.

Mice were euthanized by CO_2_ inhalation and their blood was collected immediately by cardiac puncture into EDTA-coated tubes (Sarstedt) for haematology or into lithium heparin-coated tubes (Sarstedt) for plasma biochemistry (plasma separation was performed by centrifugation at 2350g for 10 mins at room temperature, within 2 hours post-harvest). Mouse weights and liver weights were recorded post mortem. The caudate lobe of the liver was immediately frozen in liquid nitrogen, the left median lobe was frozen on dry ice and the rest of the liver was fixed for 24 hours in 10% neutral buffered formalin (in PBS), then changed to 70% ethanol before embedding.

As these are observational studies, power calculations were not routinely performed; however, animal numbers were chosen to reflect the expected magnitude of response taking into account the variability observed in pilot experiments and previous experience transcriptomic analyses. For all experiments the number of biological replicates ≥ 3 mice per cohort.

### Haematology and plasma biochemistry analysis

Whole blood haematology was performed using an IDEXX ProCyte Dx analyzer on whole blood collected in EDTA-coated tubes (Sarstedt). Biochemical analysis of plasma was carried out using a Siemens Dimension Xpand Clinical Chemistry Analyzer following International Federation of Clinical Chemistry (IFCC) approved methods.

### Histology

4μm tissue sections underwent antigen retrieval and then sequentially incubated with the primary and secondary antibody. Detection was performed with 3,3’-Diaminobenzidine and the sections were counterstained with Haematoxylin Z. Details about the antibodies and reagents can be found in Fig. S5.

Images were obtained on a Zeiss Axiovert 200 microscope using a Zeiss Axiocam MRc camera. For image analysis, stained slides were scanned using a Leica Aperio AT2 slide scanner (Leica Microsystems, UK) at 20x magnification. Quantification of blinded stained histologic sections was performed using the HALO image analysis software (V3.1.1076.363, Indica Labs). All of the slides were stained for a specific antibody in the same batch and processed at the same time in an autostainer, strictly keeping all incubation times (including that of DAB development) the same for all the samples.

For multiplex immunofluorescence, 4μm liver sections were retrieved for 25 minutes in Citrate buffer (pH 6) and were incubated with antibodies against GFP (Abcam, ab13970, 1:500), RFP (Rockland, 600-401-379, 1:200) and HNF4a (Santa Cruz, sc6556, 1:40) overnight at 4 °C. This was followed by incubation with the secondary antibodies and DAPI (1μg/μl, 0100-20, SouthernBiotech) for 1 hour at room temperature. Images were obtained using a Zeiss 710 upright confocal Z6008 microscope.

### RNA extraction

RNA extraction was performed using the Qiagen RNeasy kit (74104, Qiagen UK) as per the manufacturer’s instructions, including the optional DNase I step. Snap frozen caudate lobe (20-30mg) was homogenized using the Precellys Evolution homogenizer (Cat. Number P000062-PEVO0-A, “MET” programme) in 600μl buffer RLT/1% β-mercaptoethanol in Precellys lysing kit tubes CK14 (Precellys, P000912-LYSKO-A.0). The RNA was eluted in 30μl RNase-free water. RNA integrity and concentration were confirmed by agarose gel electrophoresis and by using the Nanodrop 2000 (Thermo Fisher Scientific) respectively. All samples had a 260/280 ratio ≥ 2.

### Quantitative reverse transcription PCR (RT-qPCR)

For RT-qPCR, RNA was extracted as described above. cDNA was generated from 1μg of RNA using the Qiagen QuantiTect Reverse transcription Kit (205313, Qiagen UK) on a PTC-200 thermal cycler (MJ Research) according to the manufacturer’s instructions. Omission of Reverse Transcriptase and a template-free reaction were used as negative controls. Quantitative real time PCR was performed with the SYBR Green system (204145, Qiagen UK) and using primers from Qiagen targeting *Per1* (QT00113337), *Per3* (QT00133455) or *Wee1* (QT00157696) using a QuantStudio 5 Real time PCR system (Thermo Fisher Scientific, A28140) in a 384 well plate setting (final reaction volume 10μl per well). Each biological replicate (mouse) was run in triplicate and 18S ribosomal RNA (Rn18S, Qiagen, QT02448075) was used as a house keeping gene for normalization.

### RNA-seq analysis

Purified RNA was tested on an Agilent 2200 TapeStation (D1000 screentape) using RNA screentape and samples with a RIN value greater than 7 were further processed for library preparation. RNA at a concentration of 20ng/μl (1μg RNA in 50μl RNase-free water) was used to prepare libraries using the TruSeq Stranded mRNA Kit. Agilent 2200 Tapestation was used to check the quality of the libraries and Qubit (Thermo Fisher Scientific) was used to assess library quantity. The libraries were then run on the Illumina NextSeq 500 using the High Output 75 cycles kit (single end, 1×75 cycle, dual index).

Raw BCL files were converted to FASTQ files using bcl2fastq2-v2.19.1 and were aligned to the mouse genome (GRCm38) using Hisat2 (v 2.1.0) and raw counts were generated using featureCounts and the GRCm38 Gencode annotation v 84. Differential gene expression was performed using edgeR. All RNA-seq analysis graphs were generated using standard R packages. Gene ontology was performed using g:Profiler (Raudvere *et al*.,2019). The raw data can be found on the Gene Expression Omnibus (GEO) repository: GSE165651.

### Statistical analyses

Statistical analyses were performed using the Prism 9 Software (GraphPad Software, Inc.). The Shapiro-Wilk test was used to assess whether data were normally distributed. For normally distributed data, either One-way ANOVA, 2-way ANOVA or the Brown-Forsythe and Welch ANOVA test was used. The Kruskal-Wallis test was performed for non parametric data. All figures were created using the Scribus Software (v1.4.7, G.N.U. general public licence). All data points on graphs represent biological replicates (each data point represents one mouse), bars represent mean ± Standard Error of Mean (S.E.M.) and P values are: *P < 0.05; **P < 0.01; ***P < 0.001 and ****P < 0.0001.

## Author contributions

CK contributed to the conceptualisation of the project, designed and performed animal studies, performed experiments, analysed data, made the figures and wrote the manuscript (original draft and subsequent editing). AW assisted with the methodology of the animal studies, performed the curation and analysis of the transcriptomics data and contributed to figure generation. CN performed the immunohistochemistry stainings. WC performed the RNA sequencing (RNA-seq). SM performed animal experiments. TGB contributed to the conceptualisation of the project, assisted with data analysis and figure generation, edited the original draft, provided resources and acquired funding. All authors contributed to the drafting and editing of the manuscript.

## Acknowledgements

CK, AW, CN, WC and SM were funded by Cancer Research UK (Grant number: A17196). TGB was funded by the Wellcome Trust (Grant number: WT107492Z). We would like to thank the CRUK Beatson Institute’s histological services, biological services and molecular technology and bioinformatics services, central services as well as the Veterinary Clinical Pathology Lab (University of Glasgow) for their help.

